# Modeling and Prediction of Body Segment Inertial Properties of Sheep from Tomographic Imaging

**DOI:** 10.1101/2025.04.30.651518

**Authors:** Aaron Henry, Carson Benner, Bailee CoVan, Annabelle Helin, Dana Gaddy, Larry J. Suva, Andrew B. Robbins

## Abstract

Estimation of body segment inertial properties (BSIPs) is a crucial step in development of inverse dynamics models. The goal of this study was to develop predictive models to estimate the mass, center of mass, and inertia tensor of the hindlimbs of sheep using easily obtainable morphometric data. In addition, this study presents a more comprehensive and repeatable method for defining each hindlimb body segment when calculating BSIPs from CT data. CT scans from 16 sheep of varying age, weight, sex, and phenotype were used to develop predictive models to estimate the BSIPs of the pelvis, thigh, crus, metatarsus, and pastern segments. The predictive models developed enable investigators to create inverse dynamics models of sheep hindlimbs. These models are particularly informative and expand the use of ovine models of human musculoskeletal disease.

## Introduction

Large animals (horses, dogs, pigs, sheep) are frequently used to study musculoskeletal diseases and orthopedic procedures, and the biomechanical impacts they have on gait and other biomechanical parameters (Khumsap et al., 2002; Brown et al., 2020; Taylor et al., 2006; Wilson et al., 2017; Thorup et al., 2008). The results of these studies are often translated into applications for human health; for example understanding the biomechanical effects of osteoarthritis or procedures such as meniscectomies (Ghosh et al., 1993; Cake et al., 2008; Moreau et al., 2014). The biomechanical techniques and models used in these animal studies must enable accurate estimation of biomechanical endpoints relevant to the disease or condition of interest, but these techniques are underdeveloped compared to those used in humans. Better biomechanical models for large animals would increase the value of biomechanical measurements from large animal studies.

Inverse dynamics (ID) is a technique used by biomechanists to calculate intersegmental reaction forces and moments from kinematic data in humans and large animals. ID models can vary in complexity and be constructed in various ways by including different numbers of body segments and restricting the degrees of freedom between each segment. Regardless of the structure of the model, inertial and morphometric properties of each segment must be specified (Manter, 1938; Seireg and Arvikar, 1973; Apkarian et al., 1989; Dogan et al., 1991; Stagni et al., 2000; Sha-har and Banks-Sills, 2002; Brown et al., 2018; Ellis et al., 2018; Brown et al., 2020). While morphometric parameters can be directly measured in living large animals (with some exceptions e.g. the hip joint center) (Henry et al., 2023), the body segment inertial properties (BSIPs); mass, center of mass, moments of inertia); cannot.

For human research, widely used predictive models often based on large datasets are used to determine subject-specific BSIPs; anthropometric data has been obtained from multiple sources including cadaver studies, balance techniques, analytical equations from geometric assumptions, stereo-photogrammetry, and medical imaging (Chandler et al., 1975; Cheng et al., 2000; Clauser et al., 1969; Crompton et al., 1996; Dumas et al., 2007; de Leva, 1996; HANAVAN, 1964; Hinrichs, 1985; Lephart, 1984; Martin et al., 1989; McConville et al., 1980; Pearsall et al., 1996; Reid, 1984; Sprigings and Leach, 1986; Young et al., 1983; Chen et al., 2011). This data is then used to develop predictive models to estimate the BSIPs. Several techniques have been used to create these predictive models including scaling equations, linear regression, and nonlinear regression (Dumas et al., 2007; de Leva, 1996; Hinrichs, 1985; Schneider and Zernicke, 1992; Yeadon and Morlock, 1989).

However, predictive models for BSIPs are not readily available for most large animals, and none of general applicability have been published for sheep. For this reason, studies investigating kinetics of use raw ground reaction force values as biomechanical endpoints, rather than intersegmental forces (Cake et al., 2008; Herfat et al., 2011; Kim and Breur, 2008; Thorup et al., 2007). This is a clear gap in our understanding of large animal kinetics and is especially important given the growing use of ovine models of human disease (Whitelaw et al., 2016; Suva et al., 2020; Perisse et al., 2021). The few studies using BSIPs for large animals typically sacrifice the animal; thus severely inhibiting the feasibility of acquiring large sample sizes and limiting the longitudinal utility of the animal (Buchner et al., 1997; Colborne et al., 2005; Duda et al., 1997; Jones et al., 2018; Nielsen et al., 2003; Taylor et al., 2006). Past investigations estimating BSIPs in sheep sacrificed the animals or used 2D imaging in the estimation, which can diminish the accuracy of the BSIPs (Duda et al., 1997; Lerner et al., 2015; Taylor et al., 2006, 2011). Furthermore, these studies have been limited by the number of sheep, often between 1 and 3, preventing investigators from creating BSIP predictive models, and therefore limiting the utility of the calculated BSIPs beyond the specific studies in which they were presented (Lerner et al., 2015; Taylor et al., 2006).

Recently, investigators have begun using medical imaging, such as computed tomography (CT) and magnetic resonance imaging (MRI), to estimate BSIPs of segments of interest in animals including birds, dogs, and even dinosaurs (Amit et al., 2009; Brown et al., 2020; Helms et al., 2009; Hutchinson et al., 2007; Paxton et al., 2014; Ragetly et al., 2008). Using imaging data, investigators calculate BSIPs by assuming tissue mass densities. However, many of the studies using this method have only calculated a singular moment of inertia and some lack predictive equations or generalizable values that could be utilized with similar large animals of varying size. Furthermore, the body segment definitions in these investigations are often unclear, difficult to replicate, and prone to labeler error. The goal of this work was to develop additional tools and methods that can be used to investigate large animal biomechanics (sheep in particular) and to mitigate the above concerns in two ways; 1) by developing predictive models to estimate BSIPs for the hindlimbs of sheep using easily obtainable morphometric data; and 2) to present a more comprehensive, reproducible method for defining each body segment when calculating BSIPs. This work is specific to sheep, but the techniques and methods developed can be directly applied to other large animals and humans.

## Methods

### Subjects and CT Scans

Sixteen Rambouillet sheep had full body CT scans (Siemens Biograph mCT) taken at 2, 7, or 14 months. Many sheep had CT scans taken at multiple time points. The present study was part of a larger investigation involving the effects of hypophosphatasia (HPP) on gait (Williams et al., 2018), and utilized existing sheep and imaging data that had previously been collected for that study. Thirty-two total CT scans from healthy wildtype (WT) (n = 9) and HPP (n = 7) sheep were used in this study.

### Anatomical Landmarking and Segment Definitions

CT data was imported into the open source software 3D Slicer and the hindlimbs were cropped 3 lumbar vertebrae cranially from the tuber coxae of the pelvis. Testes were manually removed from the CT data of male sheep and bone and skin geometries were segmented to assist in identification of anatomical landmarks and division of body segments. Trabecular and cortical bone geometries were segmented using the threshold tool at >226 Hounsfield Units (HU). Skin surface geometry was segmented at -400 HU and hollowed to a thickness of 1 mm. Voxel data external to the skin geometry was set to -1024 HU such that only tissue data would be considered in BSIP calculations. Sheep with scans at 14 months of age did not entirely fit within the CT scanner as the forelimbs and hindlimbs had to be extended. This resulted in noise around the pastern and distal metatarsus segments for fully grown sheep. This noise was manually removed, however, noise was still present within the pastern and distal metatarsus segments of these older sheep.

Virtual markers were placed on specific anatomical landmarks of both the segmented bone and skin geometries by 3 independent labelers (Table A.1). Marker locations were chosen based on ruminant dissection manuals(May, 1970; Mansour, 2017). Markers were averaged across the 3 labelers and the averaged virtual markers were used to define planes separating adjacent body segments and create body segment coordinate systems (BSCSs). These planes were created using a least squares best fit of specific groups of markers (Table A.2 and Figure A.1). Additional markers were then placed on the corners of each cut plane such that a convex hull could be created between two cut planes that encompassed a particular body segment. The thigh and pelvis segments were encompassed with convex hulls created from unique sets of markers due to their complex geometry (Table A.3, Figure A.2).

These convex hulls were then converted to binary labelmap segmentations such that each voxel within a convex hull could be attributed to a particular body segment. Joint smoothing was used between the pelvis and thigh convex hulls with priority given to the thigh segments to ensure voxels were not counted for both the pelvis and thigh. Custom Python scripts were created to automate the creation of cut planes, convex hulls, and binary labelmaps to expedite the definition of body segments within each CT scan. Specific tissues were then segmented within each body segment and used to calculate BSIPs.

### Tissue Segmentation and Calculation of Inertial Properties

Each voxel in a body segment convex hull was assigned a tissue type based on the Hounsfield unit of the voxel. Four major tissues were segmented within each body segment including cortical bone, trabecular bone, muscle, and fat. The cortical bone Hounsfield unit threshold was set to >662, trabecular bone between 226 and 661, muscle between -69 and 225, and fat between -205 and -70 (Brown et al., 2020). The 3D arrays containing the segmented tissues of each body segment were exported from 3D Slicer and processed using custom Python scripts.

A 3D mass array for each body segment was constructed using the segmented tissues of each body segment and voxel volume. The mass of each voxel was calculated as the voxel volume times the average density of the associated tissue. Voxel volume was determined from the CT settings for each scan. Average tissue density values used were based on available steer data (Gong et al., 1964) as follows: cortical bone (2.003 g/cm^3^), trabecular bone (1.911 g/cm^3^), muscle (1.06 g/cm^3^), and fat (0.95 g/cm^3^). The 3D mass array was then used to calculate BSIPs including the segment mass, center of mass, and the inertia tensor about the center of mass within the inertial frame of the CT scan.

Similar to Brown *et al.*, body segment masses were calculated as

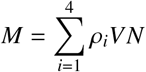

where *i* is the tissue type (cortical bone, trabecular bone, muscle, or fat), *ρ* is the tissue density, *V* is the voxel volume, and *N* is the number of voxels in the body segment (Brown et al., 2020).

The center of mass (COM) of each body segment was then calculated using the 3D mass array as follows:

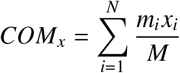

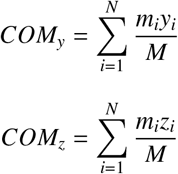

where *N* is the number of voxels in the body segment, *m_i_* is the mass of the *ith* voxel, *x_i_, y_i_*, and *z_i_* are the x,y, and z positions of the *ith* voxel, and *M* is the total mass of the segment.

The inertia tensor of each body segment was calculated as follows:

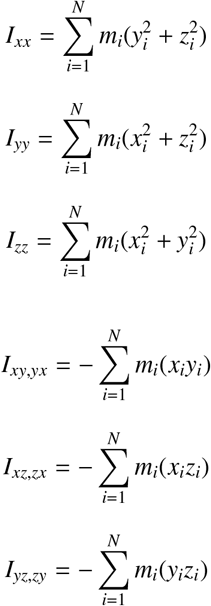

where *N* is the number of voxels in the body segment, *x_i_, y_i_*, and *z_i_* are the components of the vector between the *ith* voxel and the center of mass of the body segment, and *m_i_* is the mass of the *ith* voxel.

Segment lengths were calculated as the distance between the proximal and distal joint centers of a body segment for all segments except the pastern and pelvis. The pastern width, measured between the medial and lateral fourth metatarsal, was used for the pastern segment length. Pelvis length was measured from the midpoint of the TC markers and the midpoint of the ISC markers. Hindlimb joint widths, including the stifle, hock, and metatarsal joints, were also calculated. Stifle width was measured as the distance between the medial and lateral epicondyles of the tibia. Hock width was measured as the distance between the medial and lateral malleoli. Metatarsal width was measured as the distance between the medial and lateral fourth metatarsus.

### Body Segment Coordinate Systems and Transformation of Inertial Properties

BSIPs were transformed into each body segment’s coordinate system (BSCS) for use in motion biomechanical analysis (Tables A.4 - A.8). Segment coordinate systems were defined for each body segment using anatomical landmarks that were previously placed on the CT data. BSIPs were transformed from the inertial frame of the CT scan to the coordinate system of each body segment using the transformation below.

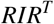

where *R* is the BSCS transformation matrix from the CT inertial frame to the body segment frame and *I* is the segment inertia tensor about the center of mass.

### BSIP Predictive Models

Several different parameters must be predicted to construct an ID model. These include the mass, the center of mass (x,y,z), and the complete inertial tensor (6 unique components) for each segment. Since left-right symmetry was assumed, this required 10 total predictive equations for each segment of the hindlimbs. Various theoretical foundations have been used in the literature to create these inertia tensor predictive equations for different populations including scaling equations (constant-only linear regression) (Dumas et al., 2007; de Leva, 1996; Amit et al., 2009), linear regression (Ragetly et al., 2008), polynomial linear regression (Buchner et al., 1997), and nonlinear regression (Yeadon and Morlock, 1989). This study investigated both scaling equations alone and scaling equations with linear regression to predict the inertia tensor of the hindlimb segments of sheep.

### Predictive Model Data

Ten of the 16 sheep were used as training data and 6 were used for testing for all predictive models. The test set was chosen such that there was sufficient diversity in age, sex, and phenotype among the test data. Both left and right body segments were used in the development of the predictive models. Additional preprocessing of input data and post processing of output data was done when necessary for each respective prediction model outlined below.

### Mass Models

Forward stepwise linear regression models were created to estimate each segment mass. The inputs for the mass regression models included age (days), sex (M/F), phenotype (WT/HPP), segment length (mm), stifle width (mm), hock width (mm), metatarsal width (mm), and body mass (kg). Minimum corrected Akaike information criteria (AICc) was used to evaluate the forward stepwise models and determine which combination of input parameters to use (Akaike, 1974). Segment mass regression models were evaluated by the relative error of the predicted mass as a percentage of the known segment mass.

### COM Models

Forward stepwise linear regression models were created to estimate the x-y-z position of the center of mass (COM) in the BSCS frame. The x-y-z position of the COM in the BSCS was first scaled by the length of the segment. Inputs to the regression models included age (days), sex (M/F), phenotype (WT/HPP), segment length (mm), and body mass (kg). Measured mediolateral (Y) position of the COM was in the negative direction for the left crus and right thigh, metatarsus, and pastern segments. To account for the differences in direction of the BSCSs between left and right limbs, the absolute value of the mediolateral (Y) position of the center of mass was used during model training. Minimum AICc was again used to evaluate the regression models. When estimating the mediolateral (Y) COM position, predicted values of the left crus and right thigh, metatarsus, and pastern segments were multiplied by -1 after estimation. COM regression models were evaluated based on the 3D Euclidian distance error as a percentage of the respective segment length between the predicted and known COM.

### Inertia Models

The inertia tensor of each segment was first scaled by the mass of the segment and the length of the segment squared (Dumas et al., 2007; de Leva, 1996). To account for the differences in direction of the BSCSs between left and right limbs, the absolute value of the Ixy and Iyz tensor elements were used during model training. Scaling equations alone and scaling equations with forward stepwise linear regression were used to predict each of the unique, scaled inertia tensor elements. The regression model included age (days), sex (M/F), and phenotype (WT/HPP) as inputs. The predicted scaled inertia tensor element was then used in combination with the predicted segment mass and measured segment length to estimate the original inertia tensor. When rescaling the Ixy and Iyz elements after estimation the following segments were negated based on differences in the left and right coordinate systems during measurement: left thigh Ixy, left pastern Ixy, right crus Ixy, right metatarsus Ixy, left thigh Iyz, left metatarsus Iyz, right crus Iyz, right pastern Iyz. The inertia tensor predictive models were evaluated by the relative error calculated as the difference between predicted inertia tensor element and the known inertia tensor element as a percentage of the maximum moment of inertia.

## Results

Average BSIPs were successfully calculated and scaled for all 32 scans (Table 1). No significant differences (p > 0.05) were found between left and right body segments of the same scan. Scaled moments of inertia (Ixx, Iyy, Izz) were expectedly higher and had far less variation across all segments than products of inertia (Ixy, Ixz, Iyz). Variation of scaled inertia tensor elements were highest for the pastern and pelvis segments. Variation of the pastern segment BSIPs increased with the inclusion of the older sheep scans that had noise around the pastern and distal metatarsus segments.

**Table 1:** Scaled BSIPs table (Mean ± standard deviation) of all 32 training and testing data points. Each parameter (column) in the table is scaled and presented as a unitless value. Mass values for each segment are scaled to the total body mass of the animal, center of mass (COM) values are scaled to the length of the segment, and inertia tensor elements are scaled as described by Dumas *et al.* and de Leva *et al.* The metatarsal width is used as the pastern length resulting in scaled BSIPs that exceed 100%.

Scaled Body Segment Inertial Properties Table
| Average Scaled Body Segment Inertial Parameters (%) |  |  |  |  |  |  |  |  |  |  |
| --- | --- | --- | --- | --- | --- | --- | --- | --- | --- | --- |
| Segment | Mass | COMx | COMy | COMz | Ixx | Iyy | Izz | Ixy / Iyx | Ixz / Izx | Iyz / Izy |
| Pastern | 0.4±0.1 | -23.4±9.7 | -5.1±8.6 | -148.9±12.9 | 100±8.5 | 100±8.3 | 51.1±6 | 8.3±3.4 | -5.8±19.1 | 17.8±8.8 |
| Metatarsus | 0.6±0.2 | -5.7±0.7 | 0.8±0.5 | -36.8±1.5 | 34.4±0.9 | 34.8±0.9 | 8.3±0.5 | 2.4±0.7 | 9.1±0.6 | 3.4±1.0 |
| Crus | 1.0±0.2 | -2.1±1.0 | -0.3±0.4 | -40.2±1.3 | 30.6±1.1 | 30.8±1.2 | 10.0±0.5 | 2.1±0.7 | -3.4±2.4 | 3.9±1.0 |
| Thigh | 6.4±0.8 | 4.2±2.3 | 4.0±2.1 | -38.0±1.8 | 36.8±1.2 | 41.0±1.1 | 27.6±2.3 | 9.8±3.0 | -18.3±1.5 | 13.5±1.6 |
| Pelvis | 5.5±0.7 | -1.8±3.0 | 0.0±0.8 | -8.9±3.5 | 27.4±2.4 | 39.6±2.1 | 38.8±1.9 | 0.6±4.0 | -14.6±1.8 | 0.9±4.1 |

### Scaled Body Segment Inertial Properties Table

### Predictive Model: Segment Mass

Segment mass predictive models were successfully created. Absolute mean relative error between the known segment mass and predicted segment mass was less than 10% for each hindlimb segment. Predictive models tended to overestimate the mass of segments with more muscle and fat including the pelvis, thigh, and crus segments (Figure 1). The outliers of the predicted metatarsus segment mass were associated with a 2 month old, female, wildtype sheep that had particularly long metatarsi compared to sheep of a similar cohort. Phenotype was a significant parameter for estimating the thigh mass, while age was significant for all segments other than the pastern (Table 2). Body mass only appeared as a significant input for segments that contained more muscle and fat.

**Table 2:** Regression constants for each segment mass. Dashed lines indicate that the parameter was not significant in predicting the segment mass. Root mean squared error (RMSE) and adjusted *R*^2^ values are included for each segment’s predictive model.

| Regression Model Constants: Segment Mass (g) |  |  |  |  |  |  |  |  |  |  |  |
| --- | --- | --- | --- | --- | --- | --- | --- | --- | --- | --- | --- |
| Segment | Intercept | Sex (F, -M) | Phenotype (WT, -HPP) | Age (Days) | Segment Length (mm) | Stifle Width (mm) | Hock Width (mm) | Met. Width (mm) | Body Mass (kg) | RMSE (g) | R<sup>2</sup> Adj |
| Pastern | -484.92 | -6.726 | - | - | - | 5.516 | - | 8.793 | - | 15.11 | 0.948 |
| Metatarsus | -766.17 | - | - | -0.107 | 1.165 | 5.322 | 5.306 | 8.374 | - | 14.47 | 0.965 |
| Crus | -759.74 | - | - | -0.268 | - | 9.971 | 1.412 | 9.587 | 5.05 | 14.62 | 0.994 |
| Thigh | -1041.23 | - | 63.622 | 1.003 | - | 27.005 | - | - | 37.59 | 132.19 | 0.989 |
| Pelvis | -1319.79 | - | - | 2.835 | 11.876 | - | - | - | 27.26 | 192.30 | 0.980 |

**Figure 1.**
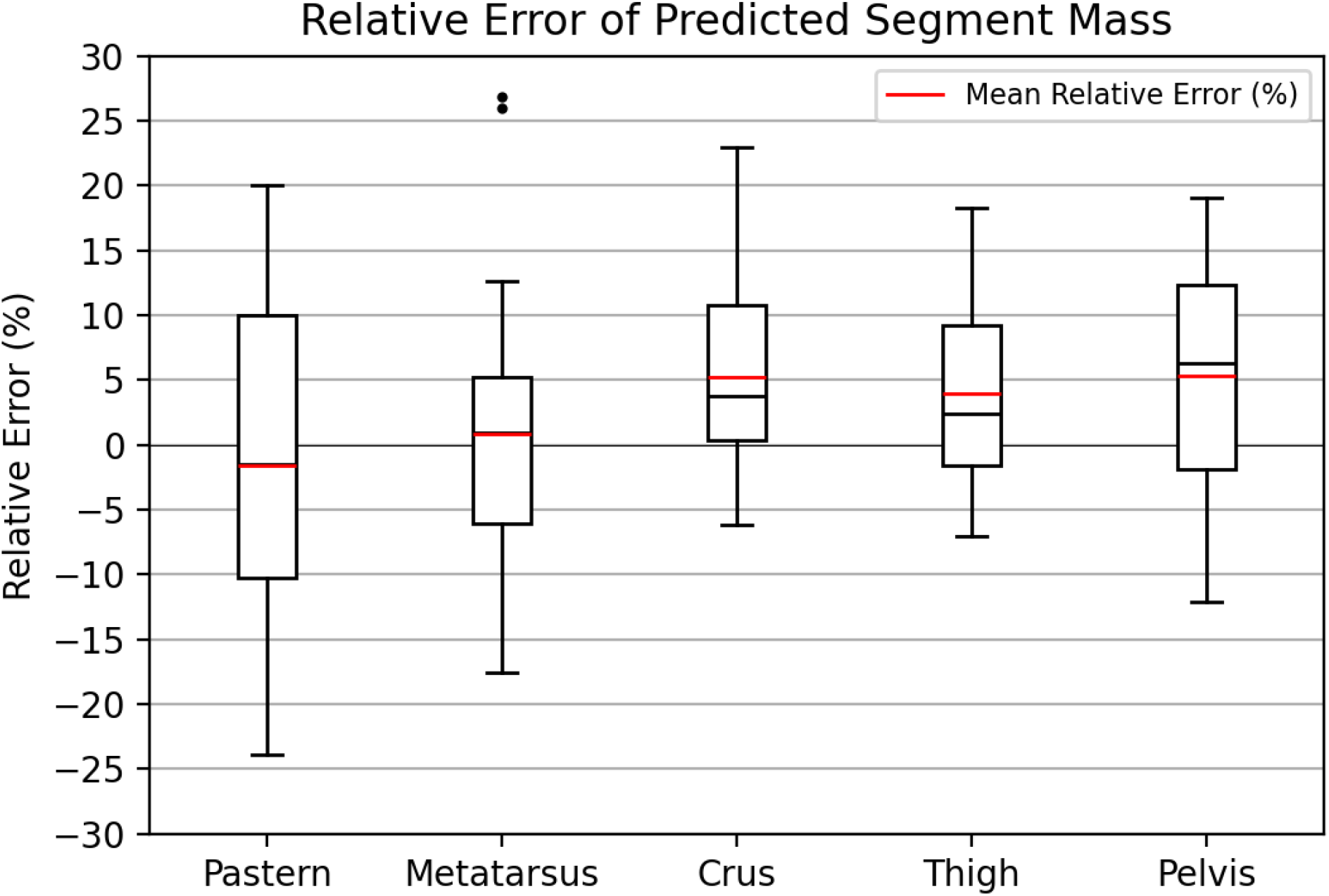
: Boxplots of the relative error of the predicted mass for each of the hindlimb segments. Relative error is calculated as the error of the predicted mass as a percentage of the known segment mass.

### Predictive Model: Center of Mass

Regression constants for each of the scaled x-y-z positions of the COM were determined for each segment (Table 3). Regression models used to predict the COM for each segment performed well apart from the pastern (Figure 2 and A.3). Mean COM error as a percentage of segment length for each segment was <10% for all segments other than the pastern.

**Table 3:** Regression constants for each coordinate of a segment’s center of mass. Dashed lines indicate that the parameter was not significant in predicting the center of mass coordinate. Root mean squared error (RMSE) and adjusted *R*^2^ values are included for each segment’s predictive model.

| Regression Model Constants: Scaled Center of Mass X (%) |  |  |  |  |  |  |  |  |
| --- | --- | --- | --- | --- | --- | --- | --- | --- |
| Segment | Intercept | Sex (F, -M) | Phenotype (WT, -HPP) | Age (Days) | Segment Length (mm) | Body Mass (kg) | RMSE (%) | R<sup>2</sup> Adj |
| Pastern | -25.322 | -1.746 | 2.336 | 0.095 | 0.313 | -0.626 | 8.21 | 0.216 |
| Metatarsus | -5.838 | - | - | - | - | - | 0.62 | 0 |
| Crus | -1.209 | - | - | - | - | -0.025 | 0.86 | 0.310 |
| Thigh | 4.082 | - | - | - | - | - | 2.12 | 0 |
| Pelvis | -3.27 | - | - | 0.008 | - | - | 2.54 | 0.172 |
| Regression Model Constants: Scaled Center of Mass Y (%) |  |  |  |  |  |  |  |  |
| Segment | Intercept | Sex (F, -M) | Phenotype (WT, -HPP) | Age (Days) | Segment Length (mm) | Body Mass (kg) | RMSE (%) | R<sup>2</sup> Adj |
| Pastern | 31.835 | - | - | - | -1.134 | 0.187 | 7.79 | 0.151 |
| Metatarsus | 0.781 | 0.112 | - | - | - | - | 0.39 | 0.056 |
| Crus | -0.214 | - | - | - | 0.004 | - | 0.30 | 0.078 |
| Thigh | 2.691 | - | - | 0.006 | - | - | 1.91 | 0.182 |
| Pelvis | 0.095 | - | - | - | - | - | 0.98 | 0 |
| Regression Model Constants: Scaled Center of Mass Z (%) |  |  |  |  |  |  |  |  |
| Segment | Intercept | Sex (F, -M) | Phenotype (WT, -HPP) | Age (Days) | Segment Length (mm) | Body Mass (kg) | RMSE (%) | R<sup>2</sup> Adj |
| Pastern | -262.322 | - | - | - | 4.736 | -0.572 | 5.05 | 0.870 |
| Metatarsus | -34.482 | - | - | - | - | -0.055 | 0.97 | 0.630 |
| Crus | -40.498 | 0.717 | -0.400 | 0.002 | - | - | 1.17 | 0.251 |
| Thigh | -37.825 | 0.622 | - | - | - | - | 1.35 | 0.163 |
| Pelvis | -32.13 | -0.219 | - | - | 0.210 | -0.241 | 2.45 | 0.372 |

**Figure 2.**
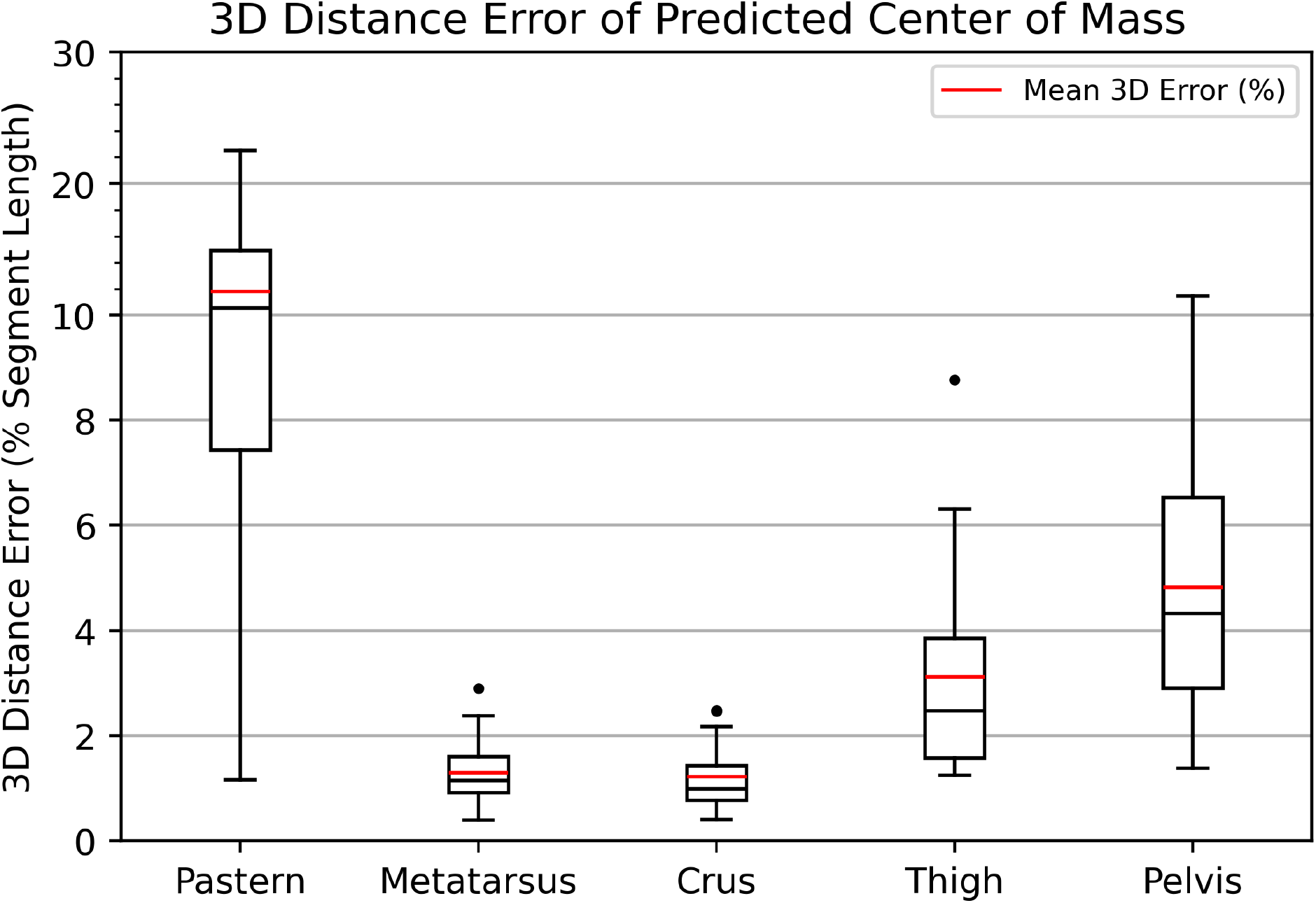
: Boxplots of the relative error of the predicted center of mass for each of the hindlimb segments. Relative error is calculated as the 3D Euclidian distance error as a percentage of the respective segment length between the predicted and known COM. The scale of the y-axis is adjusted above 10% relative error to account for the pastern segment.

### Predictive Model: Inertia Tensor

Sex and age appeared to be significant predictors sporadically across inertia tensor elements and body segments (Table 4). Phenotype only appeared to be a significant regression input for the moments of inertia of the metatarsus segment. The mean relative error of the predicted Ixx and Iyy moments of inertia appeared to be smaller for all hindlimb segments other than the pelvis (Figure 3). Relative errors for the predicted moments of inertia were higher for the pastern and metatarsus. This increase in error aligned with the increase in error of the segment mass predictive models for these segments. Products of inertia were far more difficult to predict than the moments of inertia. Relative errors were substantially higher for products of inertia, however, the relative contribution of these inertia tensor elements was small compared to the moments of inertia. Estimating the inertia tensor using only scaling equations resulted in similar performance to the linear regression approach for both moments and products of inertia (Figures A.4, A.5).

**Table 4:** Regression model constants for each inertia tensor element. When rescaling the Ixy and Iyz elements after estimation the following segments were negated to based on differences in the left and right coordinate systems during measurement: left thigh Ixy, left pastern Ixy, right crus Ixy, right metatarsus Ixy, left thigh Iyz, le1ft2metatarsus Iyz, right crus Iyz, right pastern Iyz

| Regression Model Constants: Scaled Inertia Tensor Ixx (%) |  |  |  |  |  |  |
| --- | --- | --- | --- | --- | --- | --- |
| Segment | Intercept | Sex (F, -M) | Phenotype (WT, -HPP) | Age (Days) | RMSE (%) | R<sup>2</sup> Adj |
| Pastern | 100.011 | - | - | - | 8.72 | 0 |
| Metatarsus | 34.018 | 0.261 | -0.257 | 0.004 | 0.56 | 0.566 |
| Crus | 32.104 | - | - | -0.006 | 0.73 | 0.639 |
| Thigh | 40.436 | 0.263 | - | 0.003 | 1.02 | 0.189 |
| Pelvis | 25.074 | - | - | 0.012 | 1.32 | 0.646 |
| Regression Model Constants: Scaled Inertia Tensor Iyy (%) |  |  |  |  |  |  |
| Segment | Intercept | Sex (F, -M) | Phenotype (WT, -HPP) | Age (Days) | RMSE (%) | R<sup>2</sup> Adj |
| Pastern | 100.083 | - | - | - | 9.07 | 0 |
| Metatarsus | 33.727 | 0.288 | -0.282 | 0.004 | 0.57 | 0.546 |
| Crus | 31.813 | - | - | -0.006 | 0.72 | 0.604 |
| Thigh | 37.528 | - | - | -0.003 | 1.19 | 0.145 |
| Pelvis | 38.679 | -0.869 | - | 0.005 | 1.75 | 0.220 |
| Regression Model Constants: Scaled Inertia Tensor Izz (%) |  |  |  |  |  |  |
| Segment | Intercept | Sex (F, -M) | Phenotype (WT, -HPP) | Age (Days) | RMSE (%) | R<sup>2</sup> Adj |
| Pastern | 46.496 | - | - | 0.022 | 5.33 | 0.259 |
| Metatarsus | 8.276 | -0.375 | 0.214 | - | 0.45 | 0.313 |
| Crus | 10.046 | -0.235 | - | - | 0.41 | 0.235 |
| Thigh | 25.386 | - | - | 0.011 | 1.85 | 0.443 |
| Pelvis | 39.02 | - | - | - | 1.87 | 0 |
| Regression Model Constants: Scaled Inertia Tensor Ixy / Iyx (%) |  |  |  |  |  |  |
| Segment | Intercept | Sex (F, -M) | Phenotype (WT, -HPP) | Age (Days) | RMSE (%) | R<sup>2</sup> Adj |
| Pastern | 6.041 | - | - | 0.011 | 3.41 | 0.169 |
| Metatarsus | 2.311 | - | - | - | 0.71 | 0 |
| Crus | 2.107 | - | - | - | 0.77 | 0 |
| Thigh | 9.888 | 0.998 | - | - | 3.05 | 0.080 |
| Pelvis | 0.973 | - | - | - | 3.87 | 0 |
| Regression Model Constants: Scaled Inertia Tensor Ixz / Izx (%) |  |  |  |  |  |  |
| Segment | Intercept | Sex (F, -M) | Phenotype (WT, -HPP) | Age (Days) | RMSE (%) | R<sup>2</sup> Adj |
| Pastern | 14.383 | - | - | 0.017 | 9.12 | 0.056 |
| Metatarsus | 3.494 | - | - | - | 1.10 | 0 |
| Crus | 4.563 | - | - | -0.003 | 0.93 | 0.215 |
| Thigh | 13.497 | 0.444 | - | - | 1.69 | 0.045 |
| Pelvis | -14.456 | 1.022 | - | - | 1.50 | 0.303 |
| Regression Model Constants: Scaled Inertia Tensor Iyz / Izy (%) |  |  |  |  |  |  |
| Segment | Intercept | Sex (F, -M) | Phenotype (WT, -HPP) | Age (Days) | RMSE (%) | R<sup>2</sup> Adj |
| Pastern | -18.302 | - | - | 0.062 | 17.36 | 0.208 |
| Metatarsus | 8.684 | - | - | 0.002 | 0.48 | 0.312 |
| Crus | -5.778 | - | - | 0.011 | 1.95 | 0.392 |
| Thigh | -18.15 | - | - | - | 1.51 | 0 |
| Pelvis | 0.271 | - | - | - | 3.91 | 0 |

**Figure 3.**
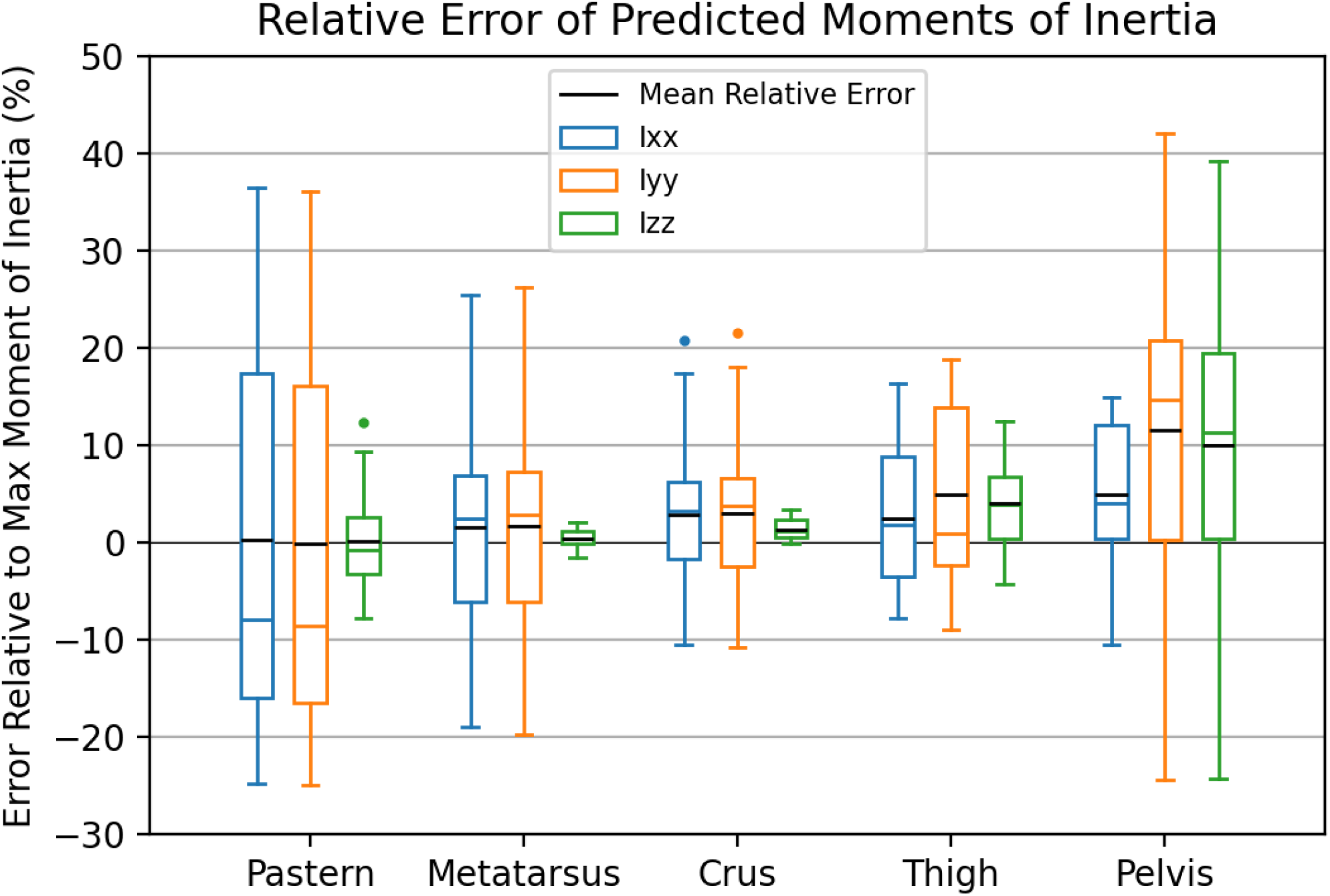
: Boxplots of the relative error of the predicted moments of inertia for each of the hindlimb segments. Relative error is calculated as the difference between predicted inertia tensor element and the known inertia tensor element as a percentage of the maximum moment of inertia.

**Figure 4.**
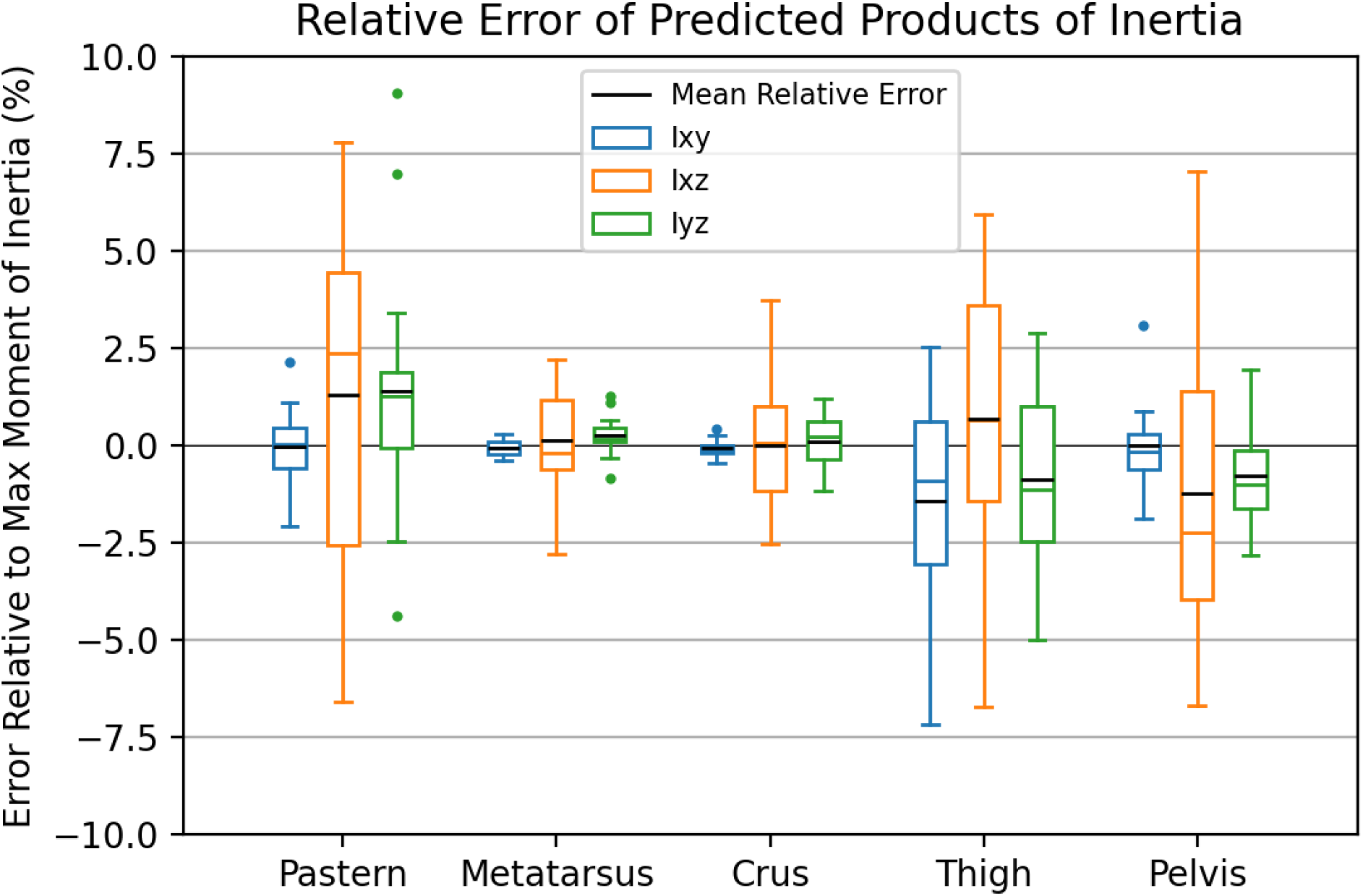
: Boxplots of the relative error of the predicted products of inertia for each of the hindlimb segments. Relative error is calculated as the difference between predicted inertia tensor element and the known inertia tensor element as a percentage of the maximum moment of inertia.

## Discussion

The goal of this study was to create predictive regression models to estimate sheep hindlimb BSIPs from CT imaging data and to provide a comprehensive and repeatable method of defining each hindlimb body segment. Placing virtual markers on CT data allowed segments to be easily distinguished from one another, segment tissues to be automatically segmented, and BSCSs and BSIPs to be calculated. Predictive regression models were successfully created to estimate BSIPs including mass, COM, and the inertia tensor for each of the hindlimb segments.

Various limitations were present in this study. Older sheep (14 months) were too large to fit entirely into the CT scanner and thus portions of the pastern and distal metatarsus had sizable amounts of noise. This noise on the exterior of the pastern and metatarsus was manually removed, although the noise inside these segments was still present. The pastern and metatarsus segments are the smallest hindlimb segments and are predominantly comprised of bone that would be less impacted by CT noise than other tissues. This noise, however, was still noticeable in the variability of BSIPs and increased error in the predicted BSIPs.

Positioning of the animal in the CT scanner is always a factor and can impact the shape of the hindlimb segments when muscles and fat are deformed. Deformation of these tissues subsequently affects the calculation of the inertia tensor as the mass distribution of a segment is altered. This was seen with variation in the pelvis BSIPs that likely arose from the whole body of the sheep shifting to one side of the scanner resulting in variation in pelvis shape segmentations. The challenge of orientation is present in all studies utilizing medical imaging of large animals as sedation is required to obtain the scans. The method used in this investigation to define body segments allows investigators to partially address this challenge with both repeatable and easily altered definitions of the cut planes used to define these body segments. Additionally, easily altered cut planes opens the possibility for researchers to investigate the impact of different cut plane locations on the resulting BSIPs that is not possible in cadaver based studies.

This study was unable to compare the calculated BSIPs to ground truth BSIPs measured with dissection methods as animals and CT images were originally acquired for a separate investigation and not originally intended for BSIP estimation (Williams et al., 2018). This study also did not evaluate the error of the regression models with data outside the range of the training data including sheep older than 14 months or of another species. Sheep are generally considered to be fully grown and mature by 12 months of age and BSIPs are not expected to be greatly altered for sheep older than 14 months, however, feeding patterns and diet may impact the size of the animal and thus the resulting BSIPs. Despite these limitations, the resulting errors of the predicted BSIPs are similar in magnitude and sometimes smaller than the limited number of prior investigations that estimated BSIPs for large animals with less than 10% mean absolute error for predicted mass, COM, and moments of inertia (Amit et al., 2009; Ragetly et al., 2008).

This investigation is the first reporting of BSIPs of sheep hindlimbs that provide predictive equations based on morphometric and demographic data and is one of the few animal studies that provides predictive equations for the entire inertia tensor for each hindlimb body segment. Furthermore, the BSIPs in this study account for the transformation of the inertia tensor into BSCS used in biomechanical analysis unlike other large animal studies (Amit et al., 2009; Ragetly et al., 2008; Buchner et al., 1997).

Various theoretical foundations for predicting the inertia tensor have been utilized in both human and large animal literature. These techniques include scaling equations (Dumas et al., 2007; Amit et al., 2009), linear regression (Ragetly et al., 2008), polynomial regression (Buchner et al., 1997), and nonlinear regression (Yeadon and Morlock, 1989). However, the associated studies were often based on a single population (age, sex) and consequently the predictive equations developed are only applicable to that specific population. The use of scaling equations and inclusion of sex, age, and phenotype in the predictive models appeared to perform the best. These additional variables were significant predictors for various inertia tensor elements with phenotype impacting the largest 3 inertia tensor elements (Ixx, Iyy, Izz) for the metatarsus only. This may be because the metatarsus is largely comprised of bone with minimal muscle and fat. The reduced density in the HPP sheep bones would suggest previously unrecognized differences for this specific segment and potential impacts on gait.

Comparable errors between the scaling only and scaling with linear regression approaches suggest that using just the scaled inertia values by themselves may be sufficient in predicting the inertia tensor for sheep hind limbs. However, the large spread of error in estimations of both approaches necessitates investigation into how these errors will propagate into calculation of joint forces and moments (Camomilla et al., 2017). The results of this study provides new information for investigators using sheep to model human musculoskeletal diseases and to investigate the kinetic impacts of medical devices. Furthermore, the improved and reproducible methodology is also easily applied to other large animal models used in the investigation of human disease, such as dogs and horses.

## Conflict of Interest

The authors declare that they have no known competing financial interests or personal relationships that could have appeared to influence the work reported in this paper.

## Ethics

This study is conducted in accordance with approved Animal Use Protocol (AUP) 2020-0164 approved by the Texas A&M University Institutional Animal Care and Use Committee (IACUC).

## Acknowledgment

This project was supported by NIH-NIDCR 1R21-DE028076 to D. Gaddy.

## Appendix

### Anatomical Landmark Markers and Joint Cut Plane Definitions

**Table A.1:**
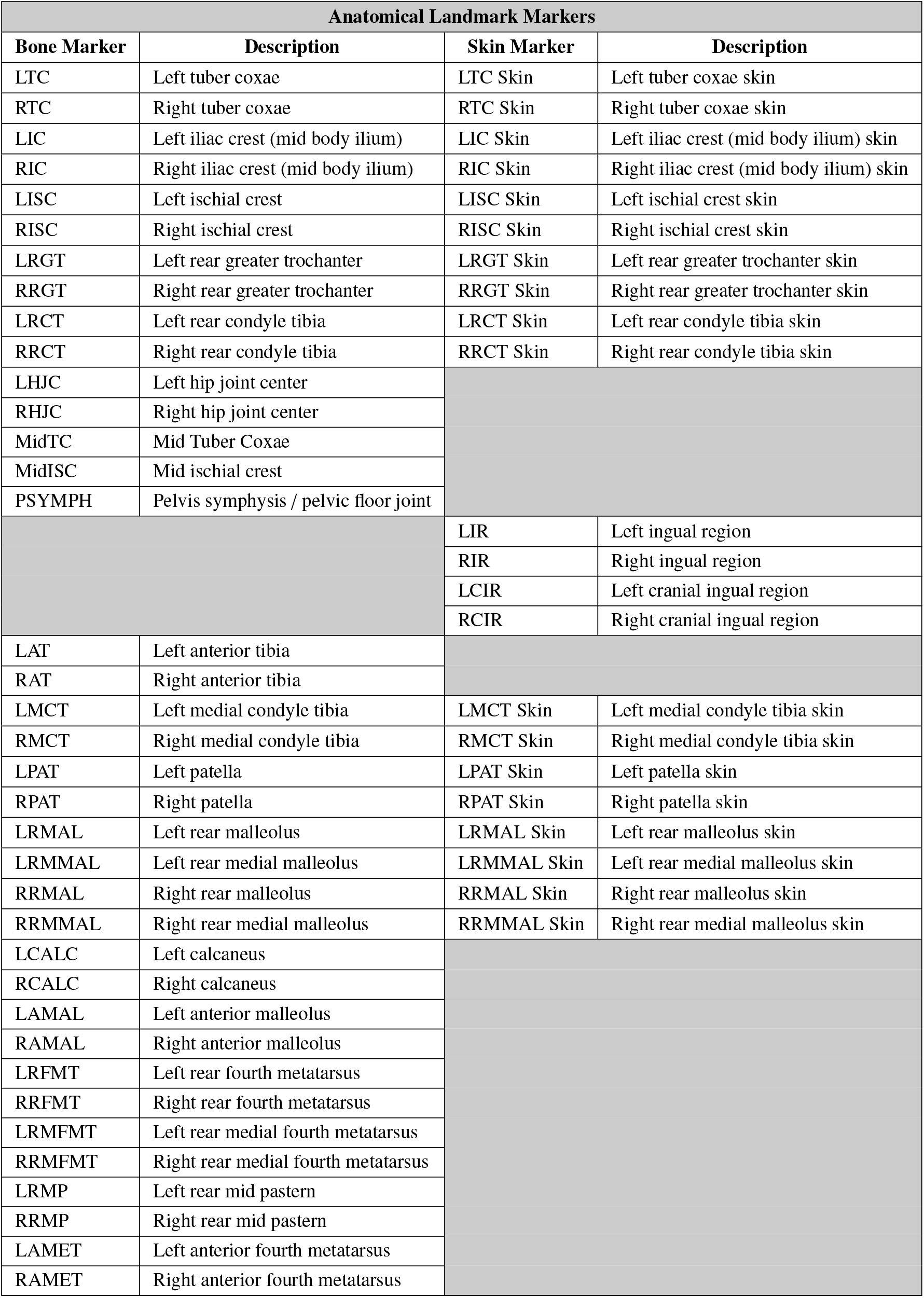
Anatomical landmark marker abbreviations and locations.

**Table A2:** Anatomical landmarks used to define joint cut planes.

| Joint Cut Plane Marker List |  |
| --- | --- |
| Joint Cut Plane | Markers |
| Sagittal Plane (SP) | MidTC, MidISC, PSYMPH |
| Right Stifle Plane (RSTP) | RRCT, RRCT Skin, RMCT, RMCT Skin, RPAT, RPAT Skin |
| Left Stifle Plane (LSTP) | LRCT, LRCT Skin, LMCT, LMCT Skin, LPAT, LPAT Skin |
| Right Hock Plane (RHP) | RRMAL, RRMAL Skin, RRMMAL, RRMMAL Skin, RCALC |
| Left Hock Plane (LHP) | LRMAL, LRMAL Skin, LRMMAL, LRMMAL Skin, LCALC |
| Right Metatarsal Plane (RMEP) | RRFMT, RRMFMT, RAMET |
| Left Metatarsal Plane (LMEP) | LRFMT, LRMFMT, LAMET |

**Figure A.1:**
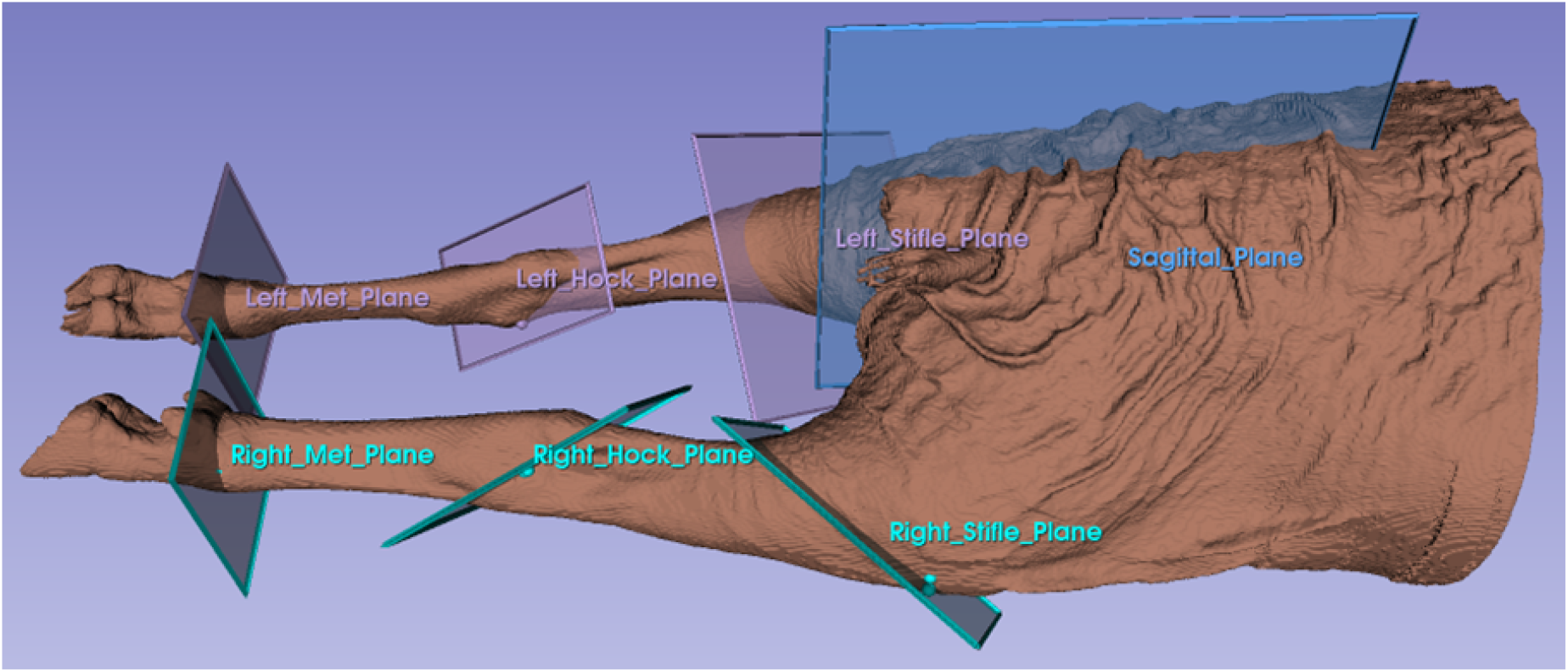
Anatomical cut planes created from Table A.2 markers.

**Table A3:**
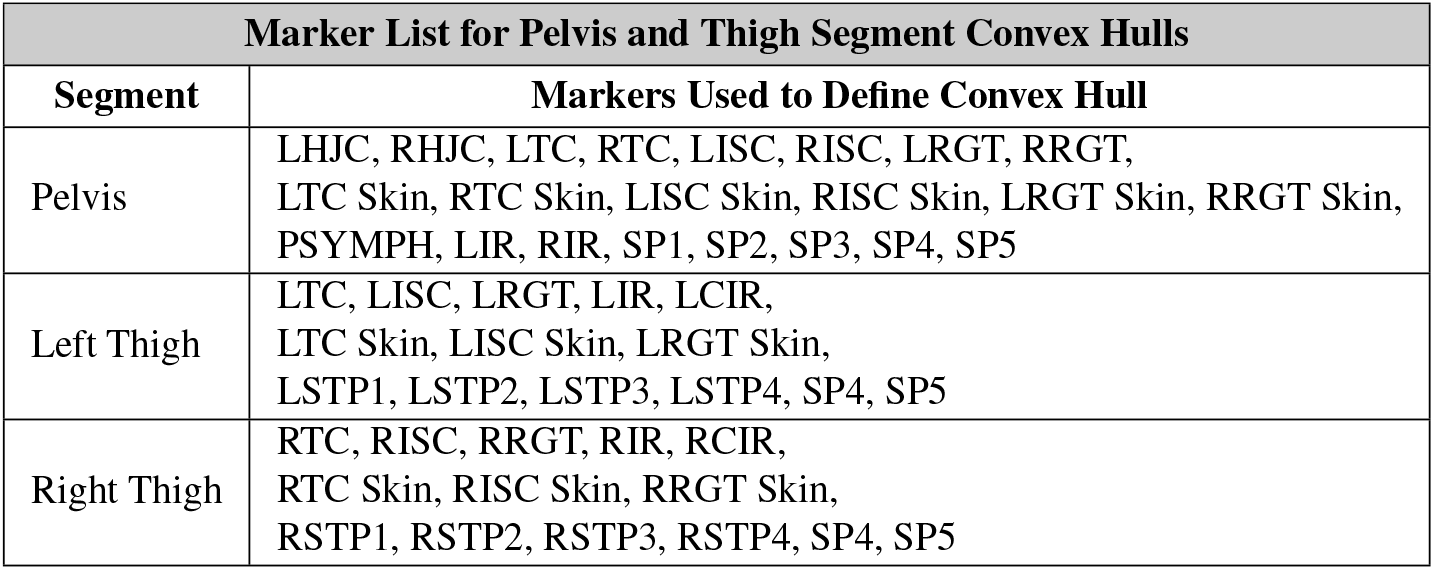
Markers used to define the convex hulls that encompass the pelvis and thigh segments. The pelvis and thigh segments use these additional markers due to their complex geometry and inability to be easily encompassed by connecting two joint cut planes.

**Figure A.2:**
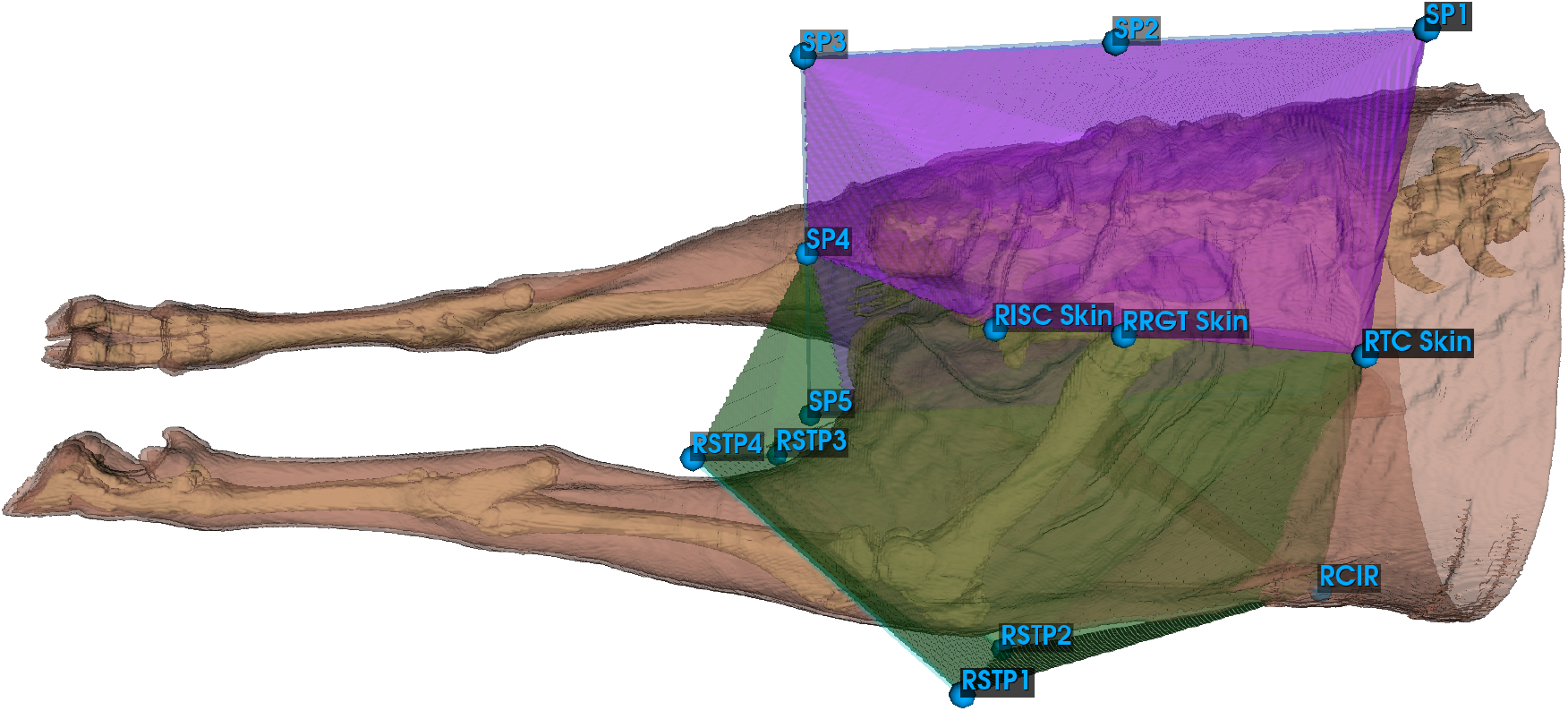
Convex hulls for the pelvis (purple) and thigh (green) created from marker lists in Table A.3. Note that not all markers from Table A.3 are visible in this figure. Sagittal plane (SP) markers were not placed on the 4 corners of the sagittal plane, rather dorsally and caudally as to extend the pelvis convex hull to encompass all tissues within the pelvis region.

### Body Segment Coordinate System (BSCS) Definitions

**Table A.4:** Definition of the pelvic body segment coordinate system.

| Pelvis Coordinate System |  |
| --- | --- |
| Origin/Axis | Definition |
| Origin | The midpoint of the LTC, RTC, LISC, and RISC skin markers. |
| X | The line connecting the midpoint of the LTC and RTC skin markers and the midpoint of the LISC and RISC skin markers pointing cranially. |
| Z | A temporary mediolateral axis is first created between the midpoint of the LTC and LISC skin markers and the midpoint of the RTC and RISC skin markers pointing to the left. The Z axis is the line perpendicular to the plane created by the cross product of the X axis and the temporary mediolateral axis, and pointing dorsally. |
| Y | The line perpendicular to the plane created by the cross product of the Z and X axes, and pointing to the left. |

**Table A.5:** Definition of the thigh body segment coordinate system.

| Thigh Coordinate System |  |
| --- | --- |
| Origin/Axis | Definition |
| Origin | The origin coincident with the left or right hip joint center. |
| Z | The line connecting the hip joint center and the stifle joint center pointing proximally. The stifle joint center is the midpoint between the medial and lateral condyles of the proximal tibia. |
| X | A temporary mediolateral axis is first created between the (L/R) MCT and (L/R) RCT skin markers pointing to the left. The X axis is the line perpendicular to the plane created by the cross product of the Z axis and the temporary mediolateral axis, and pointing cranially. |
| Y | The line perpendicular to the plane created by the cross product of the Z and X axes, and pointing to the left. |

**Table A.6:** Definition of the crus body segment coordinate system.

| Crus Coordinate System |  |
| --- | --- |
| Origin/Axis | Definition |
| Origin | The origin coincident with the left or right stifle joint center. The stifle joint center is the midpoint between the medial and lateral condyles of the proximal tibia. |
| Z | The line connecting the stifle joint center and the hock joint center pointing proximally. The hock joint center is the midpoint between the medial and lateral malleoli. |
| X | A temporary mediolateral axis is first created between the (L/R) MCT and (L/R) RCT skin markers pointing to the left. The X axis is the line perpendicular to the plane created by the cross product of the Z axis and the temporary mediolateral axis, and pointing cranially. |
| Y | The line perpendicular to the plane created by the cross product of the Z and X axes, and pointing to the left. |

**Table A.7:** Definition of the metatarsus body segment coordinate system.

| Metatarsus Coordinate System |  |
| --- | --- |
| Origin/Axis | Definition |
| Origin | The origin coincident with the left or right hock joint center. The hock joint center is the midpoint between the medial and lateral malleoli. |
| Z | The line connecting the hock joint center and the metatarsal joint center pointing proximally. The metatarsal joint center is the midpoint between the medial and lateral condyles of the fourth metatarsus. |
| X | A temporary mediolateral axis is first created between the (L/R) RMAL and (L/R) RMMAL skin markers pointing to the left. The X axis is the line perpendicular to the plane created by the cross product of the Z axis and the temporary mediolateral axis, and pointing cranially. |
| Y | The line perpendicular to the plane created by the cross product of the Z and X axes, and pointing to the left. |

**Table A.8:** Definition of the pastern body segment coordinate system.

| Pastern Coordinate System |  |
| --- | --- |
| Origin/Axis | Definition |
| Origin | The origin coincident with the left or right metatarsal joint center. The metatarsal joint center is the midpoint between the medial and lateral condyles of the fourth metatarsus. |
| Y | The line connecting the (L/R) RFMT and (L/R) RMFMT markers pointing to the left. |
| X | A temporary axis is first created between the (L/R) metatarsal joint center and (L/R) rear mid pastern marker pointing proximally. The X axis is the line perpendicular to the plane created by the cross product of the Y axis and the temporary axis, and pointing cranially. |
| Z | The line perpendicular to the plane created by the cross product of the X and Y axes, and pointing proximally. |

**Figure A.3:**
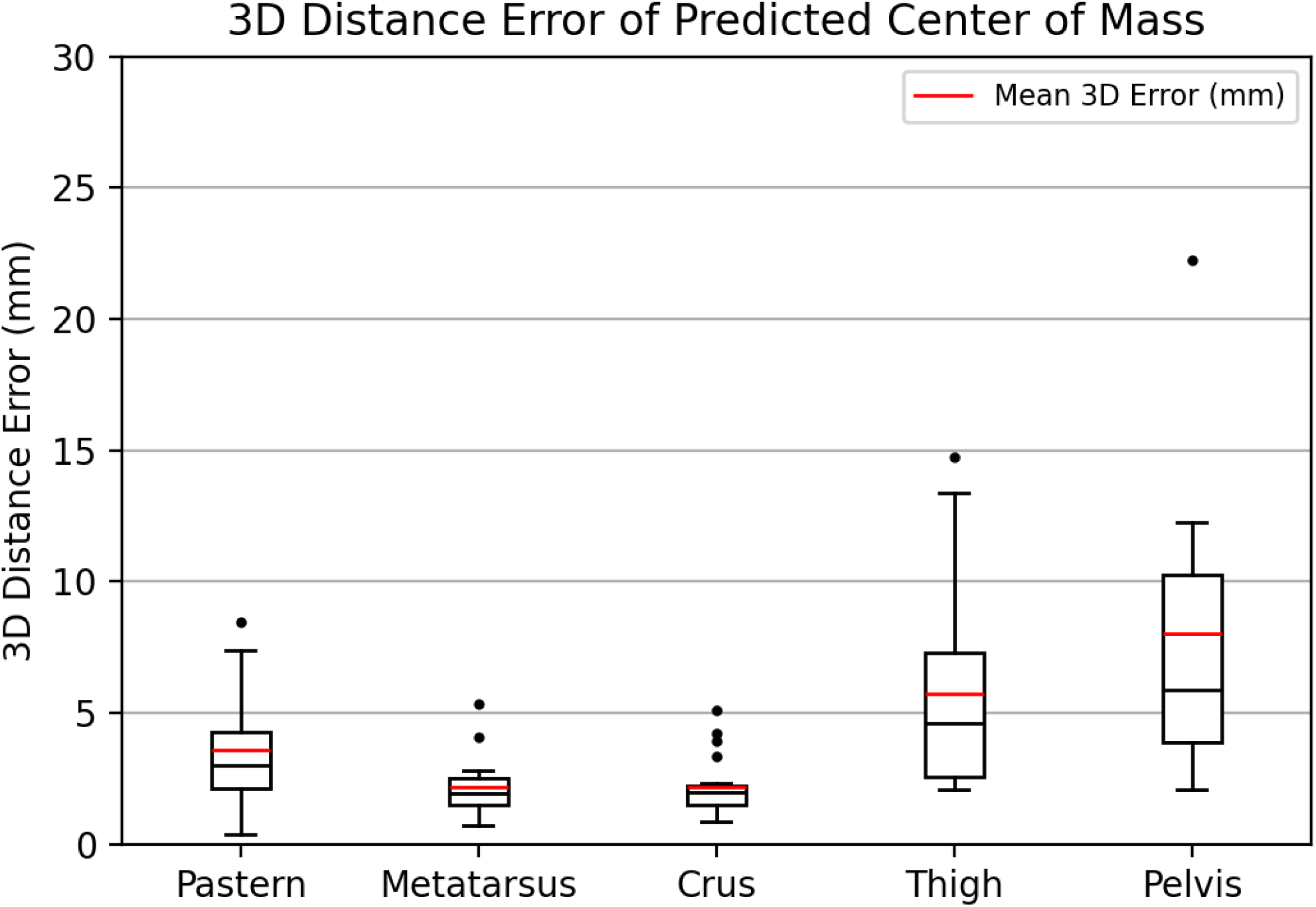
Boxplots of the absolute error of the predicted center of mass for each of the hindlimb segments. Absolute error is calculated as the 3D Euclidian distance error (mm) between the predicted and known COM.

**Figure A.4:**
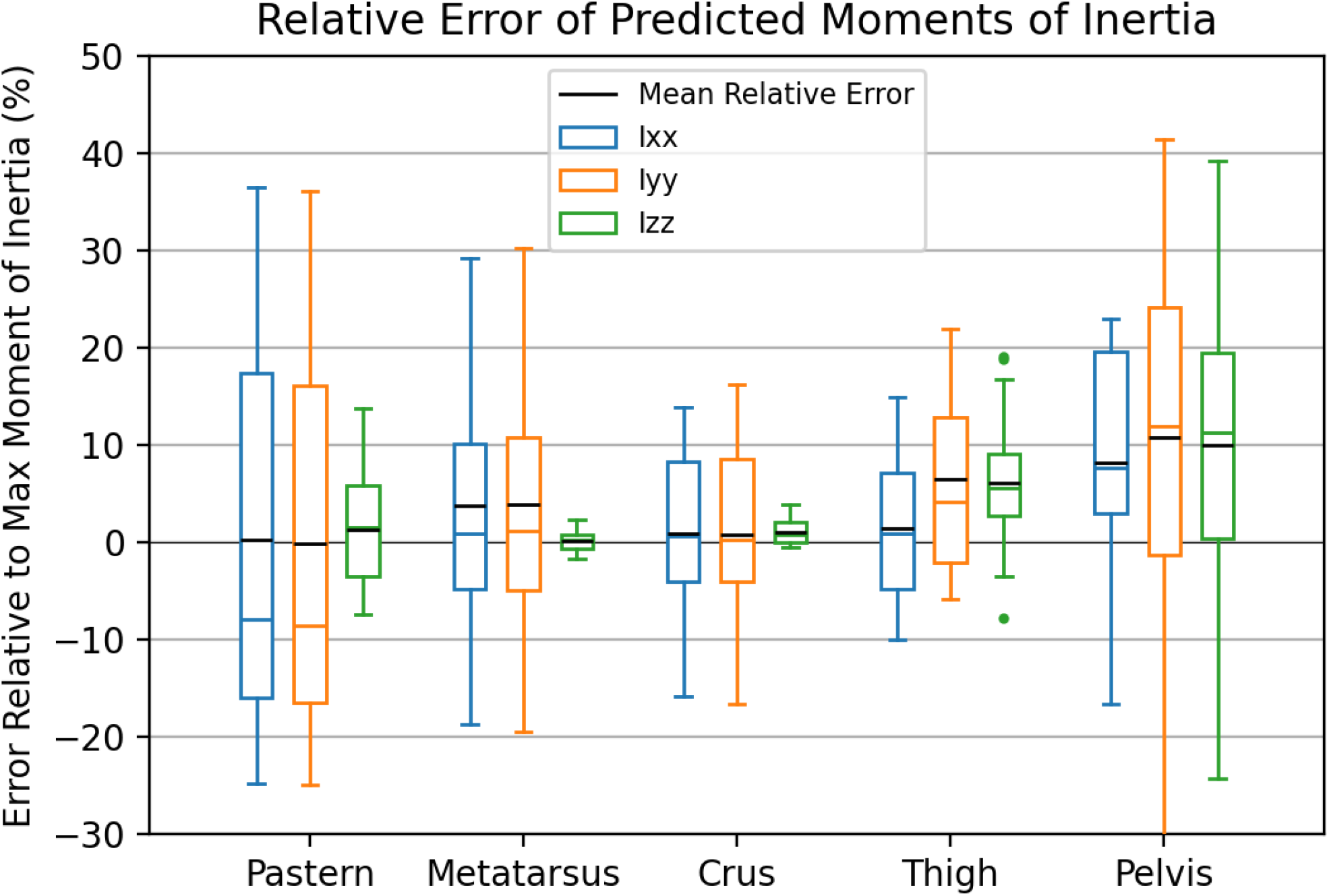
Error of predicted moments of inertia without including age, sex, or phenotype demographic data in the predictive model. Relative error is calculated as the difference between predicted inertia tensor element and the known inertia tensor element as a percentage of the maximum moment of inertia.

**Figure A5:**
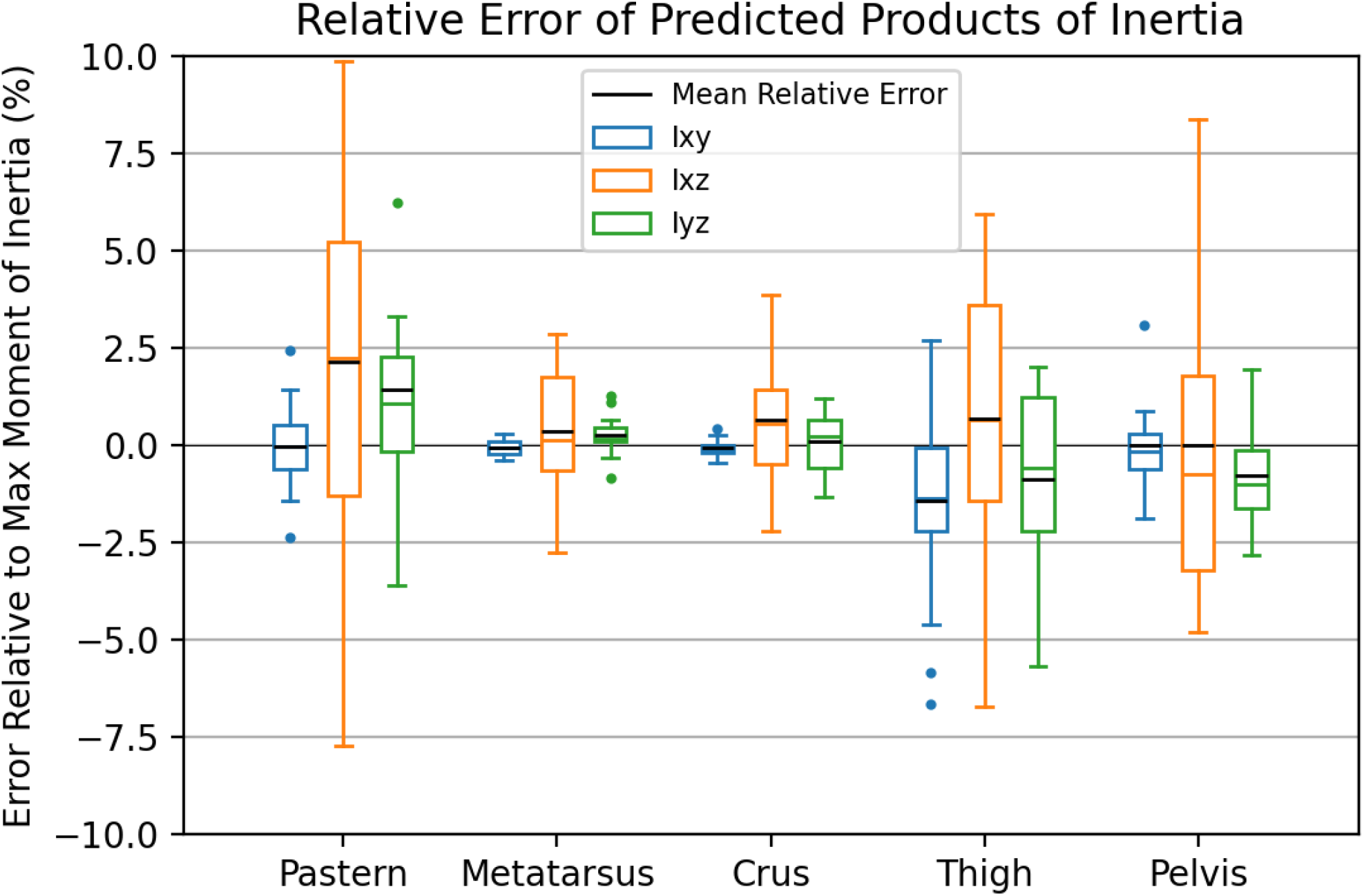
Error of predicted products of inertia without including age, sex, or phenotype demographic data in the predictive model. Relative error is calculated as the difference between predicted inertia tensor element and the known inertia tensor element as a percentage of the maximum moment of inertia.

## Notes

### Competing Interest Statement

The authors have declared no competing interest.

## References

Akaike, H. (1974). A new look at the statistical model identification. IEEE Transactions on Automatic Control, 19(6):716–723.

Amit, T., Gomberg, B. R., Milgram, J., and Shahar, R. (2009). Segmental inertial properties in dogs determined by magnetic resonance imaging. The Veterinary Journal, 182(1):94–99.

Apkarian, J., Naumann, S., and Cairns, B. (1989). A three-dimensional kinematic and dynamic model of the lower limb. Journal of Biomechanics, 22(2):143–155.

Brown, N. P., Bertocci, G. E., Cheffer, K. A., and Howland, D. R. (2018). A three dimensional multiplane kinematic model for bilateral hind limb gait analysis in cats. PLOS ONE, 13(8):e0197837.

Brown, N. P., Bertocci, G. E., States, G. J. R., Levine, G. J., Levine, J. M., and Howland, D. R. (2020). Development of a Canine Rigid Body Musculoskeletal Computer Model to Evaluate Gait. Frontiers in Bioengineering and Biotechnology, 8.

Buchner, H. H. F., Savelberg, H. H. C. M., Schamhardt, H. C., and Barneveld, A. (1997). Inertial properties of Dutch Warmblood horses. Journal of Biomechanics, 30(6):653–658.

Cake, M., Read, R., Edwards, S., Smith, M. M., Burkhardt, D., Little, C., and Ghosh, P. (2008). Changes in gait after bilateral meniscectomy in sheep: Effect of two hyaluronan preparations. Journal of Orthopaedic Science, 13(6):514–523.

Camomilla, V., Cereatti, A., Cutti, A. G., Fantozzi, S., Stagni, R., and Vannozzi, G. (2017). Methodological factors affecting joint moments estimation in clinical gait analysis: A systematic review. BioMedical Engineering OnLine, 16:106.

Chandler, R., Clauser, C., McConville, J., Reynolds, H., and Young, J. (1975). Investigation of Inertial Properties of the Human Body. Technical Report AMRL-74-137, Aerospace Medical Research Laboratory, Wright–Patterson Air Force Base, Dayton, Ohio, page 171.

Chen, S.-C., Hsieh, H.-J., Lu, T.-W., and Tseng, C.-H. (2011). A method for estimating subject-specific body segment inertial parameters in human movement analysis. Gait & Posture, 33(4):695–700.

Cheng, C.-K., Chen, H.-H., Chen, C.-S., Lee, C.-L., and Chen, C.-Y. (2000). Segment inertial properties of Chinese adults determined from magnetic resonance imaging. Clinical Biomechanics, 15(8):559–566.

Clauser, C. E., McConville, J. T., and Young, J. W. (1969). WEIGHT, VOLUME, AND CENTER OF MASS OF SEGMENTS OF THE HUMAN BODY:. Technical report, Defense Technical Information Center, Fort Belvoir, VA.

Colborne, G. R., Innes, J. F., Comerford, E. J., Owen, M. R., and Fuller, C. J. (2005). Distribution of power across the hind limb joints in Labrador Retrievers and Greyhounds. American Journal of Veterinary Research, 66(9):1563–1571.

Crompton, R. H., Li, Y., Alexander, R. McN., Wang, W., and Gunther, M. M. (1996). Segment inertial properties of primates: New techniques for laboratory and field studies of locomotion. American Journal of Physical Anthropology, 99(4):547–570.

de Leva, P. (1996). Adjustments to Zatsiorsky-Seluyanov’s segment inertia parameters. Journal of Biomechanics, 29(9):1223–1230.

Dogan, S., Manley, P. A., Vanderby, R., Kohles, S. S., Hartman, L. M., and McBeath, A. A. (1991). Canine intersegmental hip joint forces and moments before and after cemented total hip replacement. Journal of Biomechanics, 24(6):397–407.

Duda, G. N., Eckert-Hübner, K., Sokiranski, R., Kreutner, A., Miller, R., and Claes, L. (1997). Analysis of inter-fragmentary movement as a function of musculoskeletal loading conditions in sheep. Journal of Biomechanics, 31(3):201–210.

Dumas, R., Chèze, L., and Verriest, J. P. (2007). Adjustments to McConville et al. and Young et al. body segment inertial parameters. Journal of Biomechanics, 40(3):543–553.

Ellis, R. G., Rankin, J. W., and Hutchinson, J. R. (2018). Limb Kinematics, Kinetics and Muscle Dynamics During the Sit-to-Stand Transition in Greyhounds. Frontiers in Bioengineering and Biotechnology, 6.

Ghosh, P., Read, R., Armstrong, S., Wilson, D., Marshall, R., and McNair, P. (1993). The effects of intraarticular administration of hyaluronan in a model of early osteoarthritis in sheep I. Gait analysis and radiological and morphological studies. Seminars in Arthritis and Rheumatism, 22(6, Supplement 1):18–30.

Gong, J. K., Arnold, J. S., and Cohn, S. H. (1964). Composition of trabecular and cortical bone. The Anatomical Record, 149(3):325–331.

Hanavan, E. P. J. (1964). A MATHEMATICAL MODEL OF THE HUMAN BODY. AMRL-TR-64-102. AMRL-TR. Aerospace Medical Research Laboratories (U.S.), pages 1–149. Place: United States.

Helms, G., Behrens, B.-A., Stolorz, M., Wefstaedt, P., and Nolte, I. (2009). Multi-body simulation of a canine hind limb: Model development, experimental validation and calculation of ground reaction forces. BioMedical Engineering OnLine, 8(1):36.

Henry, A., Benner, C., Easwaran, A., Veerapalli, L., Gaddy, D., Suva, L. J., and Robbins, A. B. (2023). Predictive estimation of ovine hip joint centers: A regression approach. Journal of Biomechanics, 161:111861.

Herfat, S. T., Shearn, J. T., Bailey, D. L., Greiwe, R. M., Galloway, M. T., Gooch, C., and Butler, D. L. (2011). Effect of Surgery to Implant Motion and Force Sensors on Vertical Ground Reaction Forces in the Ovine Model. Journal of biomechanical engineering, 133(2):021010.

Hinrichs, R. N. (1985). Regression equations to predict segmental moments of inertia from anthropometric measurements: An extension of the data of Chandler et al. (1975). Journal of Biomechanics, 18(8):621–624.

Hutchinson, J. R., Ng-Thow-Hing, V., and Anderson, F. C. (2007). A 3D interactive method for estimating body segmental parameters in animals: Application to the turning and running performance of Tyrannosaurus rex. Journal of Theoretical Biology, 246(4):660–680.

Jones, O. Y., Raschke, S. U., and Riches, P. E. (2018). Inertial properties of the German Shepherd Dog. PLOS ONE, 13(10):e0206037.

Khumsap, S., Clayton, H. M., Lanovaz, J. L., and Bouchey, M. (2002). Effect of walking velocity on forelimb kinematics and kinetics. Equine Veterinary Journal, 34(S34):325–329.

Kim, J. and Breur, G. J. (2008). Temporospatial and kinetic characteristics of sheep walking on a pressure sensing walkway. Canadian Journal of Veterinary Research = Revue Canadienne De Recherche Veterinaire, 72(1):50–55.

Lephart, S. A. (1984). Measuring the inertial properties of cadaver segments. Journal of Biomechanics, 17(7):537–543.

Lerner, Z. F., Gadomski, B. C., Ipson, A. K., Haussler, K. K., Puttlitz, C. M., and Browning, R. C. (2015). Modulating tibiofemoral contact force in the sheep hind limb via treadmill walking: Predictions from an opensim musculoskeletal model. Journal of Orthopaedic Research, 33(8):1128–1133.

Mansour, M. (2017). Guide to Ruminant Anatomy. John Wiley & Sons, Ltd, 1 edition.

Manter, J. T. (1938). The Dynamics of Quadrupedal Walking. Journal of Experimental Biology, 15(4):522–540.

Martin, P. E., Mungiole, M., Marzke, M. W., and Longhill, J. M. (1989). The use of magnetic resonance imaging for measuring segment inertial properties. Journal of Biomechanics, 22(4):367–376.

May, N. D. S. (1970). The Anatomy of the Sheep; a Dissection Manual. [St. Lucia, Australia] University of Queensland Press.

McConville, J. T., Clauser, C. E., Churchill, T. D., Cuzzi, J., and Kaleps, I. (1980). Anthropometric Relationships of Body and Body Segment Moments of Inertia:. Technical report, Defense Technical Information Center, Fort Belvoir, VA.

Moreau, M., Lussier, B., Ballaz, L., and Troncy, E. (2014). Kinetic measurements of gait for osteoarthritis research in dogs and cats. The Canadian Veterinary Journal, 55(11):1057–1065.

Nielsen, C., Stover, S. M., Schulz, K. S., Hubbard, M., and Hawkins, D. A. (2003). Two-dimensional link-segment model of the forelimb of dogs at a walk. American Journal of Veterinary Research, 64(5):609–617.

Paxton, H., Tickle, P. G., Rankin, J. W., Codd, J. R., and Hutchinson, J. R. (2014). Anatomical and biomechanical traits of broiler chickens across ontogeny. Part II. Body segment inertial properties and muscle architecture of the pelvic limb. PeerJ, 2:e473.

Pearsall, D. J., Reid, J. G., and Livingston, L. A. (1996). Segmental inertial parameters of the human trunk as determined from computed tomography. Annals of Biomedical Engineering, 24(2):198–210.

Perisse, I. V., Fan, Z., Singina, G. N., White, K. L., and Polejaeva, I. A. (2021). Improvements in Gene Editing Technology Boost Its Applications in Livestock. Frontiers in Genetics, 11:614688.

Ragetly, C. A., Griffon, D. J., Thomas, J. E., Mostafa, A. A., Schaeffer, D. J., Pijanowski, G. J., and Hsiao-Wecksler, E. T. (2008). Noninvasive determination of body segment parameters of the hind limb in Labrador Retrievers with and without cranial cruciate ligament disease. American Journal of Veterinary Research, 69(9):1188–1196.

Reid, J. G. (1984). Physical properties of the human trunk as determined by computed tomography. Archives of Physical Medicine and Rehabilitation, 65(5):246–250.

Schneider, K. and Zernicke, R. F. (1992). Mass, center of mass, and moment of inertia estimates for infant limb segments. Journal of Biomechanics, 25(2):145–148.

Seireg, A. and Arvikar, R. (1973). A mathematical model for evaluation of forces in lower extremeties of the musculo-skeletal system. Journal of Biomechanics, 6(3):313–326.

Shahar, R. and Banks-Sills, L. (2002). Biomechanical analysis of the canine hind limb: Calculation of forces during three-legged stance. Veterinary Journal (London, England: 1997), 163(3):240–250.

Sprigings, E. and Leach, D. (1986). Standardised technique for determining the centre of gravity of body and limb segments of horses. Equine Veterinary Journal, 18(1):43–49.

Stagni, R., Leardini, A., Cappozzo, A., Grazia Benedetti, M., and Cappello, A. (2000). Effects of hip joint centre mislocation on gait analysis results. Journal of Biomechanics, 33(11):1479–1487.

Suva, L. J., Westhusin, M. E., Long, C. R., and Gaddy, D. (2020). Engineering bone phenotypes in domestic animals: Unique resources for enhancing musculoskeletal research. Bone, 130:115119.

Taylor, W. R., Ehrig, R. M., Heller, M. O., Schell, H., Seebeck, P., and Duda, G. N. (2006). Tibio-femoral joint contact forces in sheep. Journal of Biomechanics, 39(5):791–798.

Taylor, W. R., Poepplau, B. M., König, C., Ehrig, R. M., Zachow, S., Duda, G. N., and Heller, M. O. (2011). The medial–lateral force distribution in the ovine stifle joint during walking. Journal of Orthopaedic Research, 29(4):567–571.

Thorup, V. M., Laursen, B., and Jensen, B. R. (2008). Net joint kinetics in the limbs of pigs walking on concrete floor in dry and contaminated conditions1. Journal of Animal Science, 86(4):992–998.

Thorup, V. M., Tøgersen, F. Aa., Jørgensen, B., and Jensen, B. R. (2007). Biomechanical gait analysis of pigs walking on solid concrete floor. Animal, 1(5):708–715.

Whitelaw, C. B. A., Sheets, T. P., Lillico, S. G., and Telugu, B. P. (2016). Engineering large animal models of human disease. The Journal of Pathology, 238(2):247–256.

Williams, D. K., Pinzón, C., Huggins, S., Pryor, J. H., Falck, A., Herman, F., Oldeschulte, J., Chavez, M. B., Foster, B. L., White, S. H., Westhusin, M. E., Suva, L. J., Long, C. R., and Gaddy, D. (2018). Genetic engineering a large animal model of human hypophosphatasia in sheep. Scientific Reports, 8(1):16945.

Wilson, S., Abode-Iyamah, K. O., Miller, J. W., Reddy, C. G., Safayi, S., Fredericks, D. C., Jeffery, N. D., DeVries-Watson, N. A., Shivapour, S. K., Viljoen, S., Dalm, B. D., Gibson-Corley, K. N., Johnson, M. D., Gillies, G. T., and Howard, M. A. (2017). An ovine model of spinal cord injury. The Journal of Spinal Cord Medicine, 40(3):346–360.

Yeadon, M. R. and Morlock, M. (1989). The appropriate use of regression equations for the estimation of segmental inertia parameters. Journal of Biomechanics, 22(6):683–689.

Young, J. W., Chandler, R. F., Snow, C. C., Robinette, K. M., Zehner, G. F., and Loftberg, M. S. (1983). Anthropometric and mass distribution characteristics of the adult female. Technical Report FAA-AM-83-16, Civil Aeromedical Institute.

